# Rhesus macaque choices reveal that ambiguity aversion is driven by pessimistic probability estimates

**DOI:** 10.64898/2025.12.16.694644

**Authors:** Willa G. Kerkhoff, William R. Stauffer

## Abstract

Economic uncertainty has multiple forms, including risk and ambiguity. Risk describes lotteries with well defined probabilities, whereas ambiguity describes cases where probabilities are not known. Given choices between risky and ambiguous alternatives, most decision makers prefer the known uncertainty of risk. This relationship, known as ambiguity aversion, holds even when both options have the same quantitative levels of uncertainty. Despite the prevalence of ambiguity in the natural world, the psychological and neural mechanisms that produce ambiguity aversion are not well understood. Here, we developed a nonhuman primate task to study ambiguity attitudes in the lab. We showed that the animals are strongly ambiguity averse, even when they could learn the true values of the ambiguous options through repeated trials. Then, to understand why they were ambiguity averse, we examined points of subjective equivalence between risky gambles and ambiguous alternatives. These experiments demonstrated that when the potential outcomes were large, the animals judged the probability of getting the better outcome far below the true probability of 0.5. Interestingly, we observed the opposite in small value gambles: the animals were overly optimistic about their chances of getting the better reward, even though the true probability was still 0.5. These inconsistent judgments about probabilities suggest that the animals do not hold consistent beliefs, and that the lack of consistent beliefs leads to suboptimal economic outcomes. These results provide a platform to study the neurophysiological bases of real-world decision making.

## INTRODUCTION

Uncertainty is a factor in every decision because the future is unpredictable. How decision makers behave in the face of uncertainty is, therefore, a crucial component of economic decision making research. We recognise two forms of uncertainty: (1) risk, where potential outcomes and their probabilities are known, and (2) ambiguity, where outcome probabilities are unknown, undefinable, or nonexistent (Knight 1921; Mas-Colell et al. 1995). Risk is easily quantifiable but relatively rare; risk characterizes the uncertainty in coin flips and some casino games. Ambiguity is less quantifiable but much more common in everyday decisions. For example, investing in novel technologies or startups involves ambiguity, as the relevant probabilities of success are largely unknown. Decision makers tend to prefer risky over ambiguous prospects—even when the total level of uncertainty is equivalent - a phenomenon demonstrated by the Ellsberg paradox (Ellsberg 1961). This ‘ambiguity aversion’ has substantial consequences for behavior. Prices for ambiguous prospects are higher, compared to risky prospects (Bossaerts et al. 2003), investors avoid uncertain opportunities (Epstein and Schneider 2008), people skip insurance when risks are unclear (Gollier 2014), policymakers delay actions on issues like climate change (Sunstein et al. 2016), and patients often choose more conservative treatments under ambiguity (Han 2016). Thus, understanding how uncertainty influences economic behavior requires an understanding of the psychological and neural factors that drive ambiguity preferences in general and ambiguity aversion, specifically.

Ambiguous prospects necessitate subjective judgments. These subjective judgements are often interpreted within a Bayesian ‘belief’ framework (Savage 1954), but the mechanisms underlying ambiguity attitudes remain poorly understood. Prior studies have demonstrated the presence of ambiguity aversion in rhesus macaque monkeys (Hayden et al. 2010). However, it is not known what causes animals to exhibit ambiguity aversion. Here, we sought to investigate what features of choice behavior drive ambiguity aversion. We demonstrate that ambiguity attitudes are a persistent feature of NHP behavior, even in the face of learning. We further show that there is a pronounced magnitude dependence to ambiguity attitudes. Finally, we show that the magnitude dependence arises from inconsistent judgments about the probability of getting better outcomes in the face of ambiguity.

## RESULTS

All behavioral experiments were conducted in three rhesus macaque monkeys. During behavioral testing, the animals sat in a sound isolation chamber. Two of the animals made saccade-guided choices whereas a third used a touchscreen to indicate choices. We used dynamic visual cues with explicit representations of the trial-specific probabilities and magnitudes associated with each gamble (Figure 1A). To generate ambiguity, we used an occluder to hide the probability, but not the magnitude, information from the animal (Figure 1A, bottom). Each trial consisted of an initiation cue, followed by simultaneous presentation of two choice options. The animals had 2 s. to make a choice. When they selected an item, it remained on the screen while the unchosen item disappeared. The choice-dependent reward was delivered 0.5 s. later (Figure 1B).

**Figure 1.**
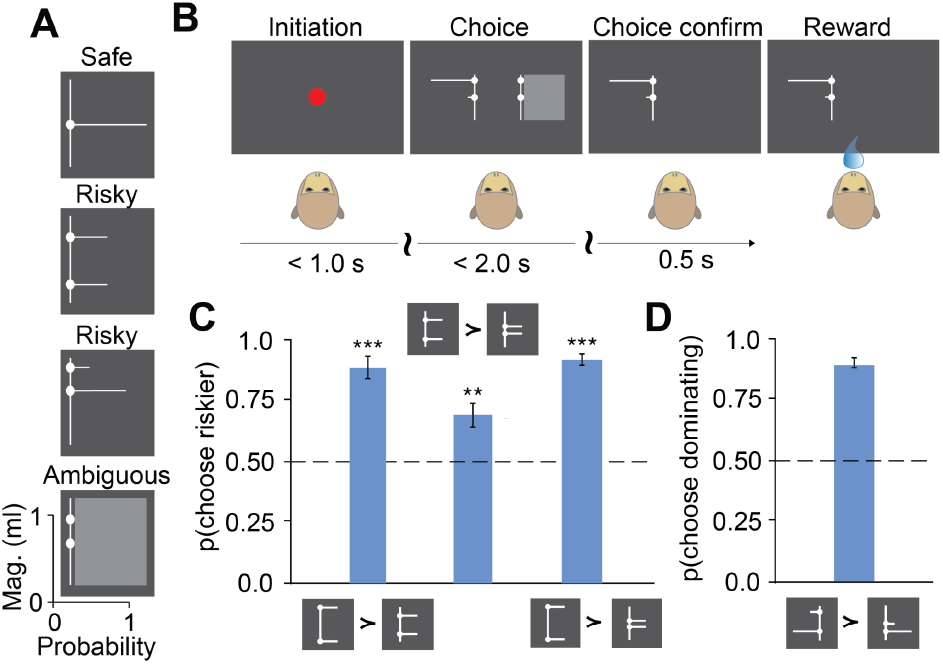
Task and basic behavior. A) Example images of visual stimuli containing explicit reward magnitude (Y-axis) and probability (X-axis) information. B) Schematic demonstrating task flow. C) Histograms indicating preferences over three pairwise choices between risky gambles that are consistent with choice transitivity. Error bars are ∓ SEM across n = 3 animals. Asterisks indicate p < 0.05 = *, < 0.01 = **, < 0.001 = ***. D) Histogram showing that the animals strongly preferred dominating to dominated options. Error bars are ∓ SEM across n = 3 animals.

### Choice behavior is consistent with basic principle of economic rationality

The animals’ behavior was consistent with basic principles of economic rationality and stochastic choice theory (Luce 1959). First, we examined choice transitivity, which states that if the animal prefers A ≻ B and B ≻ C, then the animal should prefer A ≻ C. We tested this with three gambles with equal expected values but different risks. The animals preferred the high risk gamble to the medium risk, the medium risk gamble to the low risk, and, consistent with choice transitivity, preferred the high risk gamble to the low risk gamble (Figure 1C). Thus, all animals showed a preference structure that was consistent with this basic tenant of discrete economic choice theory. Moreover, these results replicate a widely found tendency of monkeys to be risk seeking (McCoy and Platt 2005; O’Neill and Schultz 2010; O’Neill and Kobayashi 2009; Stauffer et al. 2014, 2015; Raghuraman and Padoa-Schioppa 2014; Lak et al. 2014). Next, we examined behavior under conditions where first-order stochastic dominance (FOSD) indicates the appropriate outcomes. In FOSD, the animal cannot get a worse outcome by choosing the dominating gamble, and thus should always choose it (Methods). All three monkeys chose the dominating gamble on the vast majority of trials (Figure 1D). Together, these data demonstrate that the animals’ behavior was consistent with basic assumptions of choice rationality.

### Choice behavior indicates ambiguity averse aversion

To measure uncertainty attitudes, we constructed a series of pair-wise choice problems that were inspired by the multiple price lists used in human economic experiments (Holt and Laury 2002). Our list included 5 choice pairs (Figure 2A). Both choice options were gambles, one was less risky than the other, where risk was defined as the coefficient of variance for each gamble. The choice pairs were presented repeatedly and in random order, rather than according to the list order. A risk-neutral decision maker would choose the less risky gambles in choices 1 and 2, the riskier gambles in choices 4 and 5, and be indifferent in choice 3 (Figures 2B-D, dashed line). Consistent with the transitivity test above, we observed risk seeking behavior: the animals chose the riskier option on choice pairs 3-5 more than predicted by EV maximisation (Figures 2B-D, blue plots).

**Figure 2.**
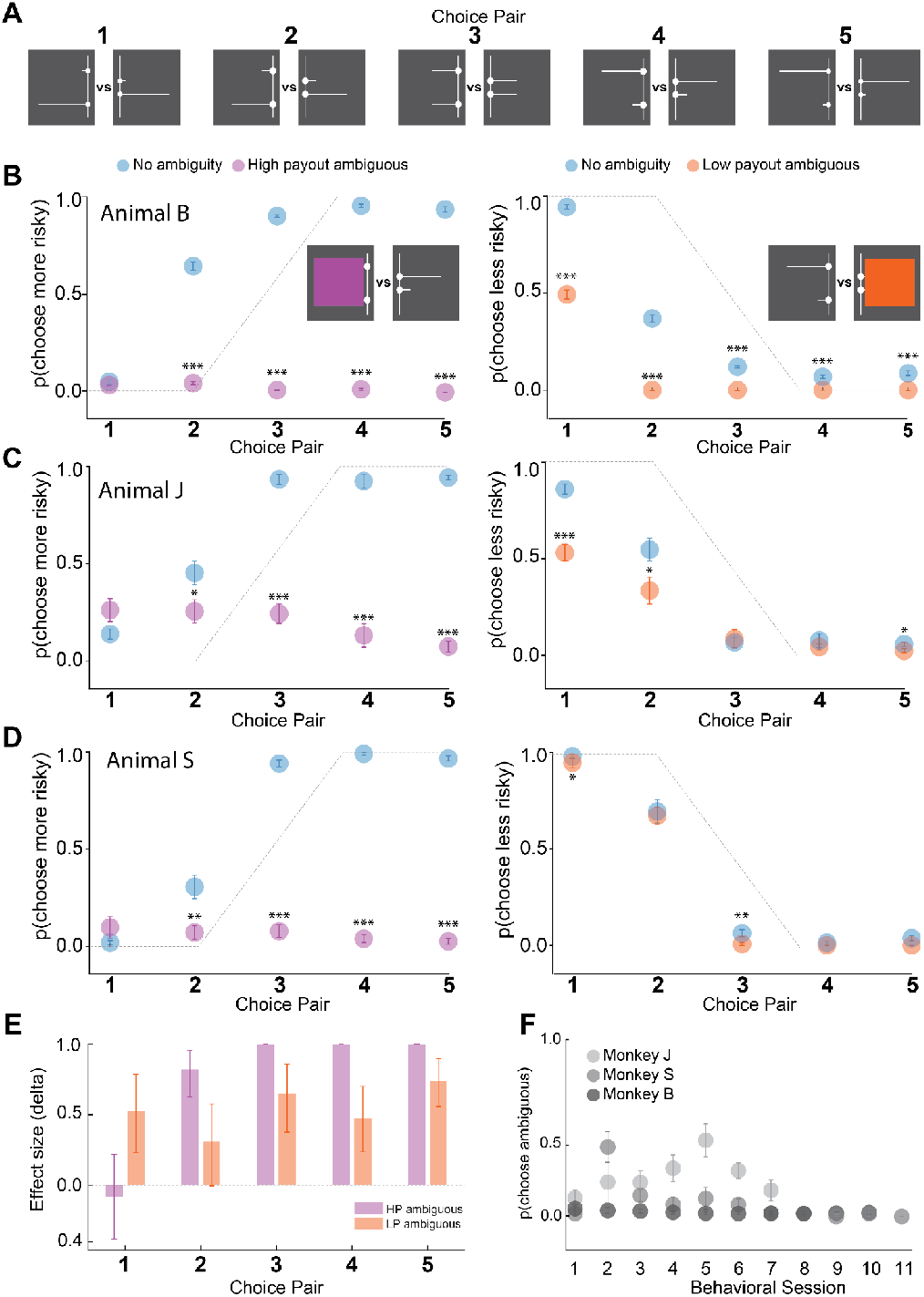
Ambiguity aversion in non-human primates. A) All choice pairs presented to animals in the multiple price list task. Each pair included a riskier stimulus (greater coefficient of variance, 1.1mL/0.1mL outcome magnitudes) and a less risky stimulus (smaller coefficient of variance, 0.85mL/0.35mL outcome magnitudes). Probabilities were matched within choice pairs (length of horizontal bars). B-D) Scatter plot of choice behavior in monkey B, J, and S respectively. Risk vs risk choices: blue, error bars ∓ SEM. Left panel: p(choose more risky) for risk only choices (blue) compared to choices where the riskier option was ambiguous (purple). Probabilities were masked with an occluder, shown in inset. Error bars are ∓ SEM, t-test risk only vs riskier ambiguous for each choice pair. Asterisks indicate p < 0.05 = *, < 0.01 = **, < 0.001 = ***. Right panel: p(choose less risky) for risky only choices (blue) compared to choices where the less risky option was ambiguous (orange). Error bars are ∓ SEM, t-test risk only vs less risky ambiguous for each choice pair. Asterisks indicate p < 0.05 = *, < 0.01 = **, < 0.001 = ***. E. Histograms show the effect sizes for ambiguity aversion when the riskier option was obscured (purple) and when the less risky option was obscured (orange). Cliff’s delta > 0 indicates ambiguity aversion, error bars are 95% CIs across animals. F) Scatter plot shows the stability of ambiguity aversion across days. Mean p(choose ambiguous) per session for each behavioral session in monkey J (lightest grey), monkey S (middle grey), and monkey B (dark grey). Error bars are ∓ SEM across trials. Spearman’s correlation showed no significant correlation between p(choose ambiguous) and session number (p > 0.05)

On a subset of trials, we made one option in each choice pair ambiguous by occluding the probabilities associated with that gamble. Note that, consistent with the definition of ambiguity, the stimuli still displayed the magnitudes (Knight 1921). On one half of ambiguity trials, we covered the probabilities associated with the riskier option (Figures 2B-D, left column), whereas on the other half of trials, we covered the probabilities associated with the less risky option.

In contrast to the risk seeking behavior we observed when we showed the animals the probabilities, we observed frank ambiguity aversion when we obscured that information. Across multiple sessions in three animals (n = 7, 10, and 11 in monkeys J, B, and S, respectively), we observed a sharp drop in the probability of choosing the riskier option when the probabilities associated with the riskier option were occluded (Figure 2B-C, left). This same ambiguity-averse behavior emerged when we occluded the less risky option - even though the animals were less likely to choose the less risky option overall (Figure 2B-C, right column). The effect sizes of the change in preferences were striking (Figure 2D, Cliff’s delta). These data demonstrate ambiguity aversion in rhesus macaques that is consistent with decades of research into human decision making.

A challenge to studying ambiguity aversion in the lab is that trials are repeated, and this provides the animals with the opportunity to learn the expected values through repeated exposures. To measure if and how ambiguity aversion persisted in the face of repetition and subsequent learning, we looked at the probability of choosing the ambiguous option as a function of session number. Monkey B showed very little shift in the probability of choosing the ambiguous option: he demonstrated nearly complete ambiguity aversion over all 10 sessions (Figure 2F). Monkeys J and S showed a minor decrease in ambiguity aversion, but the probability of choosing the ambiguous option never exceeded 50% (Figure 2C). In none of the animals did we detect a significant correlation between probability of choosing the ambiguous option and session number (p > 0.05, Spearman’s correlation). These results demonstrate that ambiguity aversion is a persistent feature of NHP behavior in this task, even when reward learning is possible.

### Ambiguity attitudes are value dependent

Our primary objective was to examine what caused ambiguity aversion. In other words, what did the animals believe was under the occluder? To measure this, we adapted the standard psychophysical method of certainty equivalents to provide us with distributional equivalents - the risky gamble that was subjectively equivalent to the ambiguous option. The method of distributional equivalents presented choices between an ambiguous option and one of five risky options with the same potential rewards. We systematically varied the probabilities of larger and smaller rewards across the five risky options: 0.1/0.9, 0.25/0.75, 0.5/0.5, 0.75/0.25, or 0.9/0.1 (Figure 3A). On each trial, the monkey made a choice via saccade or touch and received a juice reward corresponding to the selected option. We quantified behavior by computing the mean probability of choosing the ambiguous option for each risky option (Figure 3B-C). Both animals showed ambiguity aversion in the high reward range and ambiguity seeking in the low reward range. Thus, in the high value range, the animals preferred the 50-50 risky option to the high value ambiguous option, indicating pessimistic judgements about the high value ambiguous option (Figure 3B,C, yellow, p = 7.38 × 10^−4^ Monkey J, p = 0.011 Monkey S, t-test). In contrast, the animals preferred the low value ambiguous option over the 50-50 risky option with the same outcomes, indicating optimistic judgment about the low value ambiguous option (Figure 3B,C, pink, p = 9.13 × 10^−6^ Monkey J, p = 1.90 × 10^-4^ Monkey S, t-test). These results suggest that the animals beliefs about ambiguity are dependent on the value. For all three magnitude conditions, session number did not significantly predict the probability of choosing the ambiguous option (Figure 3B, logistic regression, all p > 0.05).

**Figure 3.**
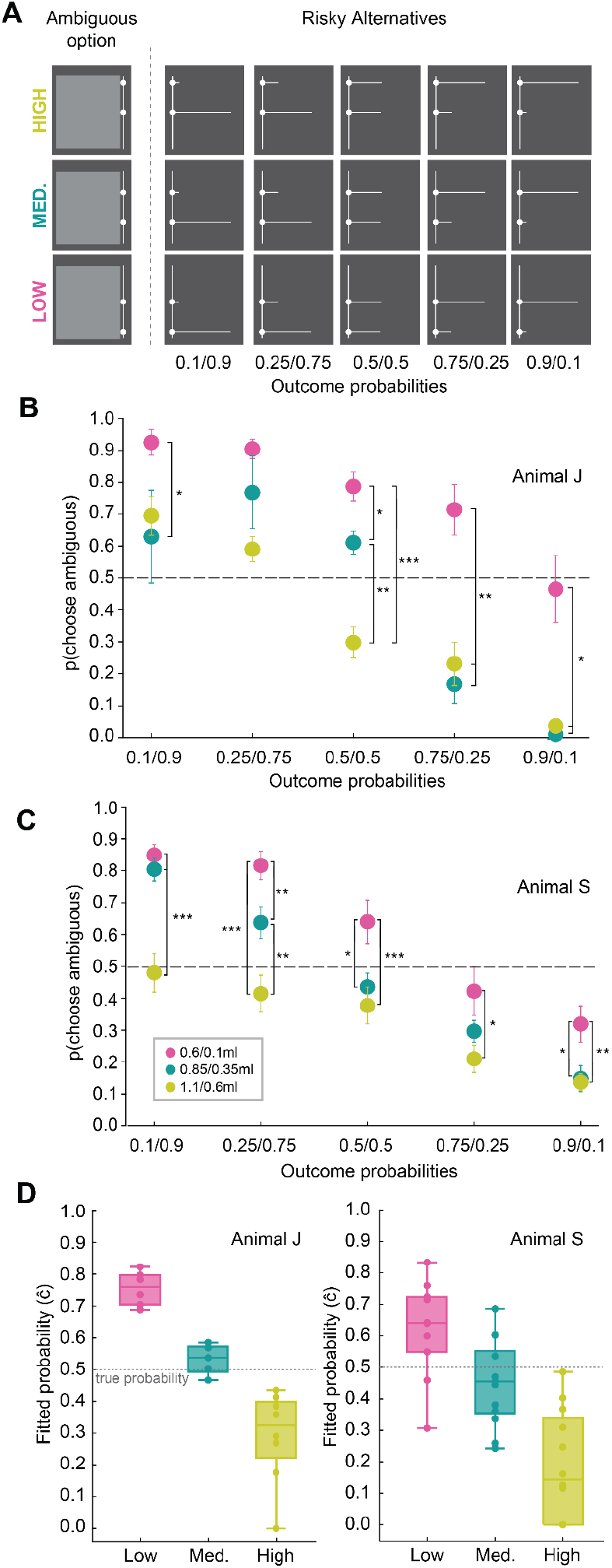
Distributional equivalents task and behavior A) Schematic showing choice pairs. Ambiguous options (left) were paired with one of five risky alternatives (right) that had the same outcome magnitudes. Magnitude conditions: High, 1.1mL/0.6mL juice outcomes, yellow. Medium, 0.85mL/0.35mL juice outcomes, green. Low, 0.6mL/0.1mL juice outcomes, pink. B) Scatter plot showing mean p(chose ambiguous) across all choice pairs and all behavioral sessions in Monkey J. C) As in B, for Monkey S. Error bars in B and C are ∓ SEM. p < 0.05 = *, < 0.01 = **, < 0.001 = ***. D) Box plots of distributional equivalents fitted per session for all magnitude conditions in Monkey J (left) and S (right). Distributional equivalents modeled as ĉ under the α-maxmin model of expected utility. Dots reflect fitted measures for each session. Box-and-whisker plots show the median, the interquartile range (box; 25th–75th percentiles), and whiskers extending to 1.5 times the interquartile range.

We fit a logistic function derived from the α-maxmin model of expected utility (Gilboa and Schmeidler 1989; Ghirardato et al. 2004) to the choice probabilities and defined the distributional equivalent as the logistic function’s midpoint (Methods). In practice, this procedure fitted the maximum likelihood probability of getting the better option, given the animals’ choices. The fitted probabilities reflected the distributional equivalents (Figure 3D): the fitted probabilities in the low value gambles were always above 0.5, whereas the fitted probabilities for the high value gambles were always below 0.5. These data demonstrate a choice inconsistency in the face of ambiguity, in a manner dependent on the reward outcomes on offer.

## DISCUSSION

Here we show that rhesus monkeys display persistent ambiguity attitudes. The animal’s choices indicated that they understood the task and that their goal was maximising reward value (Figure 1). As in previous studies, the animals were risk seeking (McCoy and Platt 2005; O’Neill and Schultz 2010; O’Neill and Kobayashi 2009; Stauffer et al. 2014, 2015; Raghuraman and Padoa-Schioppa 2014; Lak et al. 2014). In contrast to their risk seeking attitudes, they displayed frank ambiguity aversion (Figure 2). We designed a choice paradigm to find the risky gamble that was subjectively equivalent to the ambiguous prospect, which we denoted as the distributional equivalent. We found that the animals overweighted the chance of getting the better outcome in low value gambles, whereas the distributional equivalent for high-magnitude gambles indicated underweighting the chance of the better outcome. Choice models fit to the data showed that the animals behaved ‘as if’ the probability of the better outcome was close to 0.7 in small gambles, but behaved ‘as if’ the probability of the better outcome in the large gambles was very small.

Ambiguity is a crucial factor in most decisions. Because ambiguous probabilities are not, by definition, provided or even existent, decision making under ambiguity is entirely dependent on decision makers’ thoughts and emotions. Consider that, one major factor driving ambiguity aversion is ‘fear of negative evaluation’ (Curley et al. 1986). In fact, when ambiguous preferences are unknown to others, ambiguity aversion often disappears (Trautmann et al. 2008). This finding demonstrates the purely psychological nature of ambiguity attitudes. Our results provide a foundation to study the neurophysiological mechanisms that drive these psychological processes while animals make choices under ambiguity.

## ACKNOWLEDMENTS

The authors thank Rachel Tittle for reading the manuscript and providing feedback. We also thank Jacquelyn Breter for animal care. This work was supported by the National Institute of Mental Health (NIMH) with award number R01MH128669.

## METHODS

All animal procedures were approved by the Institutional Animal Care and Use Committee of the University of Pittsburgh. We used three male Rhesus macaque monkeys (*Macaca mulatta)* in these studies (Monkey J: 6 years of age, 7.85 kg. Monkey B: 12 years of age, 13.0 kg. Monkey S: 12 years of age, 10.2 kg). Under general anesthesia, all monkeys were surgically implanted with a head post (Grey Matter Research, Rogue Research) over parietal and occipital cortex. During tasks, monkeys sat in a primate chair (Crist Instruments, Rogue Research) positioned 30cm in front of a touchscreen-enabled computer monitor during experiments. Monkeys reported choices by gaze-fixating on items on the screen or by touching items. Behavioral sessions took place daily and consisted of ∼250 to 350 behavioral trials. We monitored eye movements non-invasively during experiments using infrared eye tracking (Eyelink Plus 1000). Custom-made software (MATLAB, Mathworks Inc.) was used to control the behavioral task. In the case of Monkey S, a small subset of the data presented was collected while the animal was freely moving in the home cage. An in-cage training system (Thomas RECORDING GmbH) was attached to the front of the cage and the task was presented using a touchscreen Samsung tablet. The in-cage version of the task was controlled using custom-made software (Android) running on the touchscreen tablet. In both settings, blackcurrant juice was delivered as reward via a task-controlled solenoid valve.

### BEHAVIORAL TASKS

#### Basic choice behavior

For the data shown in Figure 1, we presented monkeys with pair-wise choices between visually informative conditioned stimuli (Figure 1A). The cues predicted both the magnitude of possible rewards and the probabilities associated with those rewards. All monkeys were extensively trained before data collection to ensure that they understood the information presented in the cues. Monkeys initiated all behavioral trials (Figure 1B). Following initiation, monkeys were immediately shown two choice options and allowed to indicate their choice any time within 2 s. Following selection, the unselected choice option disappeared and reward was delivered. Trials where the monkey failed to initiate the trial or failed to make a selection within the time window resulted in an error sound, a 4 second time-out, and repeat of the trial. Monkeys learned that selecting a stimulus would result in a given reward amount being delivered with the probability indicated by the stimulus (Figure 1A). Our stimulus design allows us to flexibly offer lotteries to our animals where any reward magnitude could be paired with any probability between 0 and 1. The reward range was between 0.1mL and 1.2mL of blackcurrant juice, and probabilities always summed to 1 for a given stimulus. To demonstrate that our animals learned the stimuli parameters and used the information they gained to make rational economic decisions, we present data from two experiments where our monkeys demonstrate adherence to two key economic principles: choice transitivity and first-order stochastic dominance (FOSD).

Choice transitivity is formalized as if *A*≻*B* and *B*≻*C*⟹ *A*≻*C*. We presented our monkeys with pair-wise choices between three stimuli with the same expected value but different levels of risk (Figure 1C, example stimuli). The risk for a given choice option is defined as the coefficient of variation, calculated by dividing the standard deviation by the mean return. Stimuli with a greater reward range, and thus a larger standard deviation, have a greater coefficient of variation and are therefore riskier than stimuli with a smaller reward range. Both monkeys preferred the high risk option to the medium risk option, and the medium risk option to the low risk option (Figure 1C, blue left two histograms). Choice transitivity indicates, therefore, that the animals should choose the high over the low risk option. This is indeed what we observed (Figure 1C, right histograms). This result demonstrates that the animals choice behavior was consistent with choice transitivity.

FOSD is a key criterion that must be obeyed in order to infer that the animals are maximizing utility when choosing between two gambles. The dominance relationship between two gambles is defined by their cumulative distribution functions (CDFs) (Mas-Colell et al. 1995). If the CDF of one option is always higher, that option first-order stochastically dominates the other option. Intuitively, one choice option dominates the second choice option if nothing can be lost by selecting the first. To examine if the animals’ choices obeyed FOSD, we presented our monkeys with pair-wise choices where one option first-order stochastically dominated the other (Figure 1D, example stimuli). Animals strongly preferred the dominating option (Figure 1D, blue histogram), demonstrating that they both understood the information presented in the stimuli and that they consistently used that information to maximize utility. Together, these data demonstrate that the animals’ behavior was consistent with basic assumptions of choice rationality.

#### Choice task to demonstrate risk seeking and ambiguity aversion

To measure uncertainty attitudes, we constructed a series of pair-wise choice problems that were inspired by the multiple price lists used in economic experiments in humans (Holt and Laury 2002). Each choice pair consisted of two two-item gambles. Gambles in the same pair had identical probability distributions but different reward magnitudes. As with the choice transitivity experiments, we made one gamble in each pair riskier than the other (Figure 2A, riskier stimulus presented on left for each choice pair). Our monkeys reported their choices via saccade or touch and showed consistent risk seeking (Figure 2B-D, blue).

To measure ambiguity attitudes, we introduced a grey occluder that covered the probability information for one stimulus on a subset of trials (Figure 1A, bottom). The occluder was on the screen for every trial in a behavioral session. For the trials where we measured risk attitude, the occluder was on the screen but not covering either choice option. On trials with ambiguity, we placed the occluder over the horizontal probability bars of one stimulus but left the dots (which indicate reward magnitude) unobstructed. Thus, the animal could still determine the potential reward outcomes of the ambiguous stimulus, but the probabilities associated with those outcomes was unknown. In a behavioral session, one third of the trials an animal completed were choices between two risky stimuli. Of the remaining two thirds, half were trials where the riskier stimulus was ambiguous (Figure 2B-D, purple) and half were trials where the less risky stimulus was ambiguous (Figure 2B-D, orange). Each animal completed between 250 and 350 trials per session.

#### Choice task to reveal equivalence between risky and ambiguous options

In Figure 3, we report data from a pair-wise choice task that allows direct measurement of the subjective values of ambiguous options. Similar to the approach employed to measure risk attitudes using certainty equivalents, we had monkeys make choices between risky lotteries and ambiguous lotteries with matched outcome magnitudes (Figure 3A). We then employ an ɑ-maxmin model of expected utility (Gilboa and Schmeidler 1989) (Gilboa and Schmeidler 1989; Ghirardato et al. 2004)(Gilboa and Schmeidler 1989) to describe ambiguity attitudes across three magnitude conditions (Low, Medium, High) in two monkeys (Monkey J, S, Figure 3B-C). In this model, subject behavior is summarized by a single parameter, c, which represents the risky probability at which the subject is indifferent between the ambiguous option and a risky lottery. Under the α-maxmin framework, the c is given by:

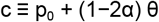

where p_0_ denotes the center of the belief set over ambiguous probabilities, θ reflects the degree of ambiguity, and α governs the relative weight placed on pessimistic versus optimistic beliefs. Lower values of c indicate ambiguity aversion, corresponding to more pessimistic evaluations of the ambiguous option, whereas higher values of c indicate ambiguity seeking, corresponding to more optimistic evaluations. Accordingly, the distribution of ĉ for each monkey serves as our primary measure of ambiguity attitudes across magnitude conditions (Figure 3D). We fit a logistic function with terms described under α-maxmin expected utility to choice data from each behavioral session across each of the three conditions in both monkeys. c was then reported per session (Figure 3D). We performed logistic regression to determine if the behavioral session was predictive of ambiguity attitude, and found no significant effect.

